# CVTree: A Parallel Alignment-free Phylogeny and Taxonomy Tool based on Composition Vectors of Genomes

**DOI:** 10.1101/2021.02.04.429726

**Authors:** Guanghong Zuo

## Abstract

CVTree is an alignment-free algorithm to infer phylogenetic relationships from genome sequences. It had been successfully applied to study phylogeny and taxonomy of viruses, prokaryotes, and fungi based on the whole genomes, as well as chloroplasts, mitochondria, and metagenomes. Here we presented the standalone software for the CVTree algorithm. In the software, a parallel workflow for the CVTree algorithm was designed. Based on the workflow, new alignment-free methods were also implemented. And by examining the phylogeny and taxonomy of 13903 prokaryotes based on 16S rRNA sequences, we showed that CVTree software is an efficient and effective tool for the studying of phylogeny and taxonomy based on genome sequences.

**Availability:** https://github.com/ghzuo/cvtree

## Introduction

Comparative analysis of genome sequences is the fundamental approach for the phylogenetic study of biology. Traditionally, sequence comparison was based on pairwise, including global [1] and local [2] sequence alignments, or multiple sequence alignment (MSA). Software tools for sequence alignment, such as BLAST [3] and CLUSTAL [4], are the most widely used bioinformatics methods. The aligned sequences provided a very intuitive impression of the difference between sequences, especially for the sequences with high identity. However, the computation of an accurate MSA is an NP-hard problem. The MSA-based methods cannot be solved in a realistic time for the large data sets that are available today. Most of MSA tools are based on heuristic algorithms. It was found that alignment-based techniques were inaccurate in scenarios of low sequence identity [5, 6]. Therefore, as an alternative solution to sequence alignment, many alignment-free approaches to sequence analysis have been developed in recent decades [6–10]. These methods are computationally less expensive than the alignment-based methods. Their scalability allows them to be applied to much larger data sets than conventional MSA-based methods.

CVTree stands for *Composition Vector Tree*, which is a cluster of alignment-free methods based on subsequences of a defined length (named as k-string). They generated dissimilarity matrices from a comparatively large collection of genome sequences for phylogenetic studies. The classical CVTree method was proposed by Prof. Bailin Hao and coworkers in 2004 [11]. In the classical CVTree algorithm, every genome sequence, including protein, RNA, and DNA, was represented by a composition vector, which was calculated by the difference between the frequencies of k-strings and the prediction frequencies by the Markov model. And the similarity between two sequences was measured by the cosine of two composition vectors. The classical CVTree method was first developed to infer evolutionary relatedness of Bacteria and Archaea [11–14], and then successfully applied to fungi [15, 16], viruses [17], chloroplasts [18], and mitochondria [19], as well as metagenomes [20, 21]. After the proposal of the classical CVTree method, there are three versions of CVTree web server were released successively by our group [22–24]. The latest released CVTree web server, CVTree3 [24] is available from http://tlife.fudan.edu.cn/cvtree (Fudan University, Shanghai) and http://bigd.big.ac.cn/cvtree (Beijing Institute of Genomics, Beijing).

In this paper, we presented the standalone software for the CVTree algorithms. Due to the flexibility of the standalone software, the CVTree software is helpful for the researchers who are interested in the intermediate results, e.g., the collection of CVs and dissimilarity matrices, or unwilling to upload their data to web server, as well as bioinformatics developers. In the CVTree software, the programs were redesigned in an object-oriented model. The OpenMP technique was employed to make the main programs parallel. An inbuilt automatic workflow helps users to obtain the phylogenetic tree from the FASTA files directly, and the intermediate result was cached to avoid redundant calculation. Based on the scheme of the CVTree algorithm, other alignment-free phylogenetic methods based on the composition vectors were implemented [25]. Furthermore, by using CVTree software, we obtained the phylogeny of 13903 prokaryotes based on their 16S rRNA sequences [26]. Interestingly, it was found that these CVTree methods have better performance than that of the alignment-based method in both efficiency and accuracy.

## Algorithms and Implementations

### Scheme of CVTree

CVTree include a cluster of alignment-free methods to obtain phylogenetic relationship based on genome sequences. Figure 1 showed the scheme of the CVTree methods. There are three steps for the algorithm, i.e., modeling the genomes to the composition vectors, calculating the dissimilarity matrix from the composition vectors, and inferring phylogenetic tree based on the dissimilarity matrix. In the CVTree software, the classical CVTree composition vector generative model and algorithm were named as Hao model and Hao method to honor Prof. Bailin Hao [27]. And in the Hao method, the genome sequences were cut into small k-strings. Then the composition vector of the genome was modeled by the frequencies of k-strings, including the length k-2, k-1, and k, based on a Markov model. The dissimilarity of two genomes was measured by the cosine of the angle between two vectors. Finally, the phylogenetic tree was inferred by the neighbor-joint algorithm. Based on the scheme, other conventional dissimilarity methods, including Jaccard, Manhattan, Euclidean, etc., were integrated into the CVTree software [25]. Two composition vector models, i.e., count and Hao model, and an enhanced tree neighbor-joint method are also provided in the software. Users can compose the models and methods by the options of programs (see detail in manual).

**Figure 1.**
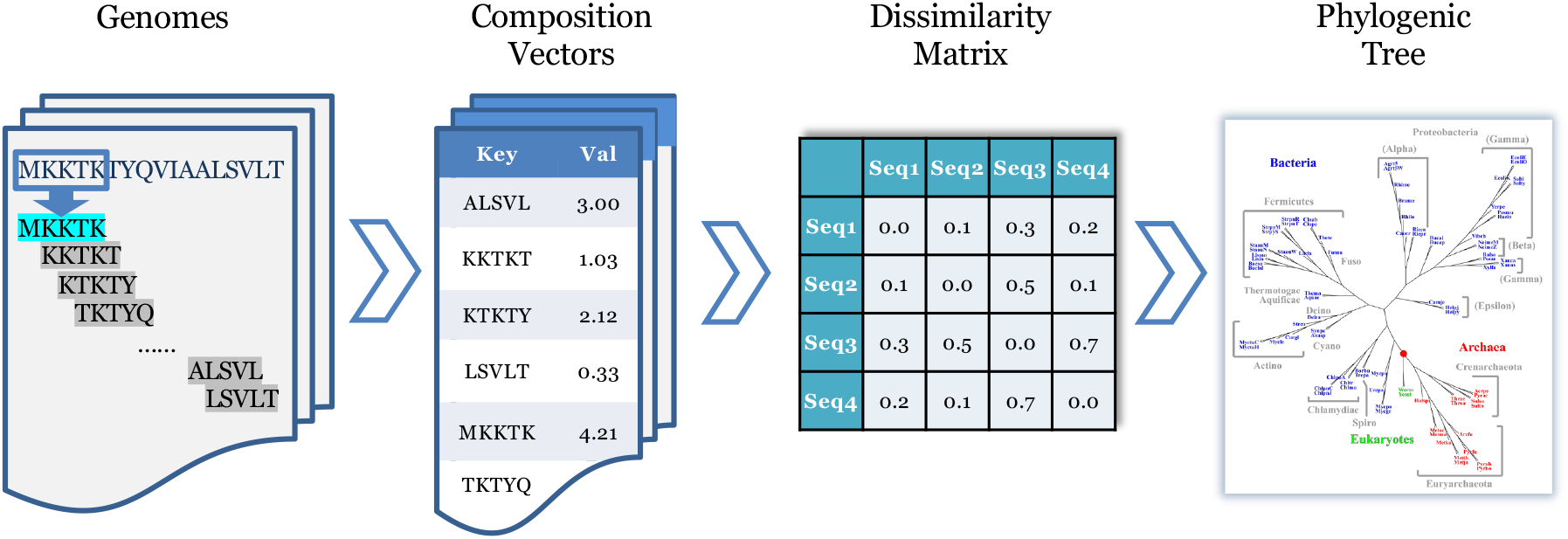
Scheme of CVTree algorithms. Four block indicated the four different states of the information, i.e., genomes, composition vectors, dissimilarity matrix, and phylogenetic tree in CVTree algorithm. And three chevrons indicated three steps to handle the information flow along these four states, i.e., from genomes to composition vectors, from composition vectors to dissimilarity matrix, and from dissimilarity matrix to phylogenetic tree, in the CVTree algorithm.

### Implementation

The CVTree software, written in standard C++, facilitates compilation compliant with CMake and execution on Linux/Unix, Macintosh, and Windows platforms. The CVTree software, including example data, documentation, and source codes, is freely available for academic use at https://github.com/ghzuo/cvtree.

The CVTree programs are object-oriented design. As the scheme of CVTree (see figure 1), there are four states for the information, i.e., genomes, composition vectors, dissimilarity matrix, and phylogenetic tree. They are described by different classes in CVTree programs. In more detail, the k-strings were encoded into an unsigned long integer (64-bit length) to improve efficiency. It is obvious that for a *N* length sequence which consisted of *m* letters, when the length *k* of k-string is large enough, the number of k-string in the sequence, *N* − *k* + 1, is much less than that of the type of k-string, *m*^*k*^. That is, the composition vector is sparse, i.e., most of dimensions are zero. Thus, only the non-zero dimensions were saved as key-value pairs in CVTree programs. All composition vectors were handled by associated arrays in the generation and by sorted sequential arrays in the calculation, respectively. The operations on these four states were also described by classes. And to organize different methods, we designed three virtual classes as the interface to describe the three operations in the CVTree scheme. In this way, a new method can be implemented by deriving from corresponded base classes (see detail in the manual). To improve efficiency, the main programs of CVTree are implemented in parallel by OpenMP techniques. And these classes were carefully designed to keep threads safe.

The input data for CVTree software are the genomes in Fasta form, in which one file contains one genome. And a file contains the list of the genomes is also required. The final output of CVTree is the phylogenic tree in Newick form. As the scheme of CVTree algorithms, there are three steps from genomes to a phylogenic tree. Thus, there are three programs, named as *g2cv, cv2dm*, and *dm2tree*, to perform these three tasks respectively. Apart from the step-by-step way, an integrated program, named as *cvtree*, is also provided in the CVTree software. Figure 2 showed the flowchart of the *cvtree* program. Instead of a bundle of those three steps programs, the *cvtree* program automatically refers to the intermediate data to reduce computing resource consumption. Therefore, apart from the final phylogenic tree, the intermediate data, including composition vectors and dissimilarity matrices, are also saved for reuse in the next calculation. And to save the storage, these intermediate data are compressed into binary format, which cannot be inspected directly. Thus, the tools to handle these compressed files were also provided in the CVTree software (see detail in manual).

**Figure 2.**
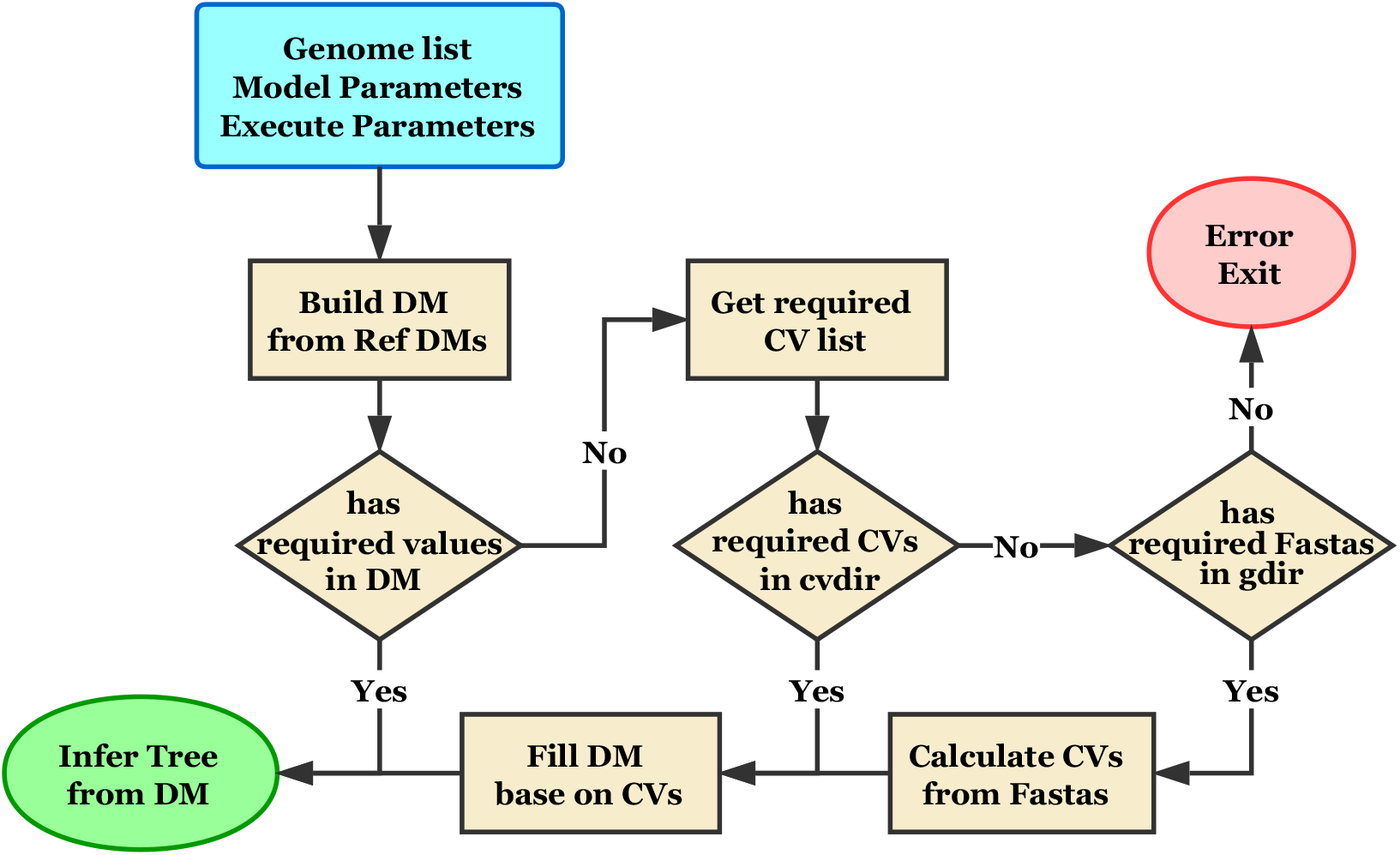
Flowchart of cvtree command. The workflow of the cvtree command was to start from the blue square, follow the arrow line, normal end at the green ellipse. In the workflow, the cvtree command automatically checks the cache data of every step to avoid redundant calculation.

## Results and Discussions

The CVTree was first developed to infer evolutionary relatedness based on whole genomes to obtain real species trees, instead of gene trees. The phylogenetic tree for prokaryotes based on whole genomes can be accessed on the CVTree3 web servers by an interactive interface. As an example, here we showed the CVTree software by performing cvtree on 13904 RNA sequences, in which 13903 sequences were 16S rRNA sequences from the LTPs132 of the “All-Species Living” project [26] and one sequence was from the virus, *Ferret parechovirus*, as the outgroup to root the phylogenetic tree. It was found that the performance of *cvtree* is very remarkable. By the acceleration of multi-core CPUs, a typical phylogenetic tree for these 13904 sequences can be obtained in 108.8 seconds on our Dell PowerEdge Server (4 × 20-Core Intel Xeon Gold 6248 @ 2.50GHz, Linux System), or in 493.4 seconds on our Apple MacBook Pro (8-Core Intel Core i9 @ 2.3 GHz, MacOS System). Figure 3 showed the elapsed time of *cvtree* as a function of the number of threads in our Dell PowerEdge Server. As figure 3, the speedup of parallel was very significant. The calculation was accelerated about 1.76 times when the number of threads doubled. Detail studies showed that most of the time was spent in the last two steps, calculating dissimilarity matrix and inferring phylogenetic tree. And the speedup by the parallel in calculating dissimilarity matrices is more significant. It was about 1.91 times with double of threads. We noted that with the increase of the length of genome sequences, the complexity of calculating dissimilarity matrices is lower than linear complexity, and that of inferring phylogenetic tree is constant. This is, the amount of computing resource of CVTree methods is scaled with the length of sequence below linear. And CVTree method may obtain a rich benefit by parallel. Therefore, the CVTree programs are efficient enough to obtain the all-species living tree based on whole genomes.

**Figure 3.**
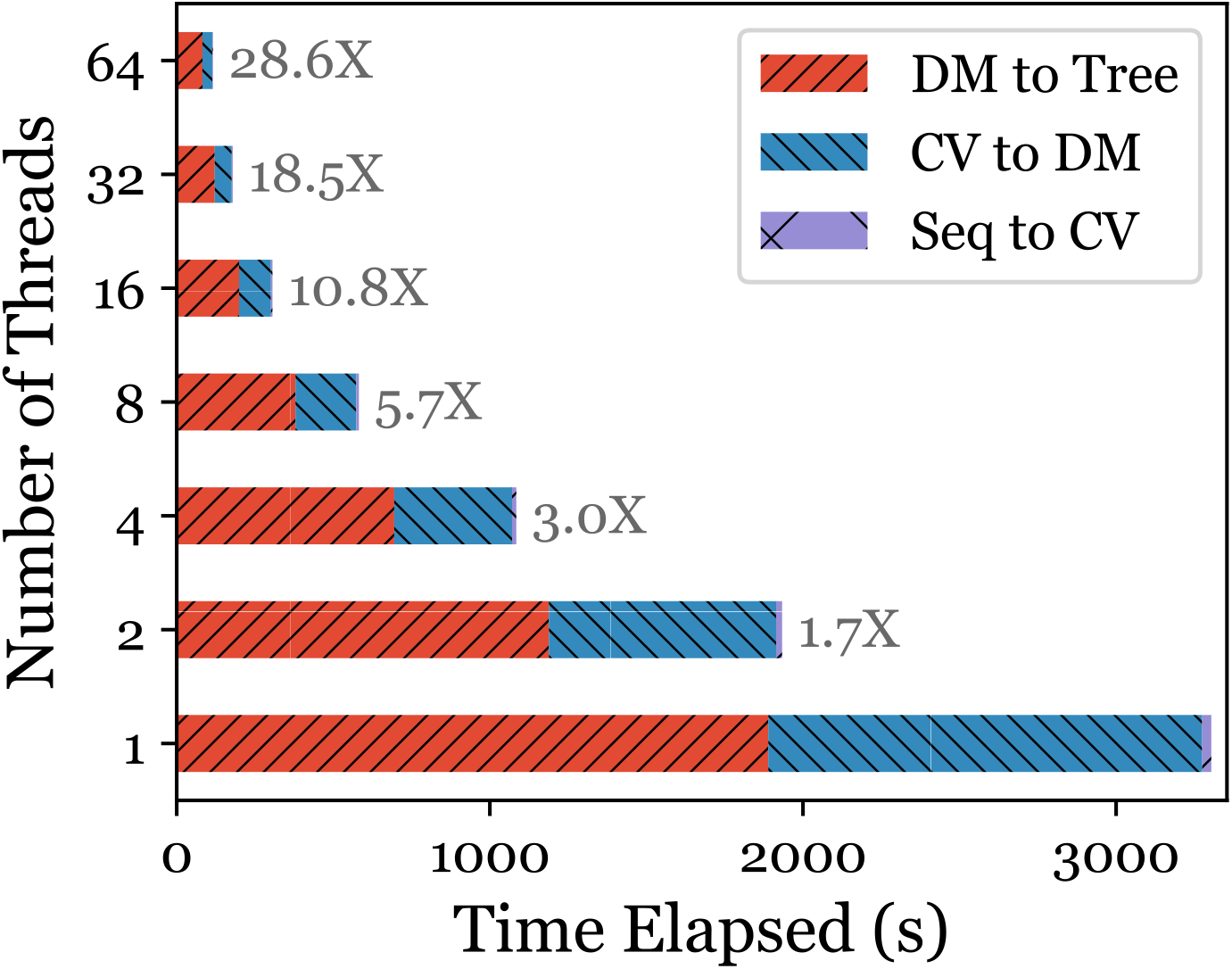
Speedup of cvtree command by multi-thread. Here the Hao model with *k* = 6 was performed by cvtree command on the DELL PowerEdge Server (4 × 20-Core Intel Xeon Gold 6248 @ 2.50GHz, Linux System). The elapsed time of the three steps displayed in different patterns and colors in the bar plot. And the number at the right of the bars showed the speedup of the cvtree command by multi-thread.

To examine the accuracy of CVTree, we compared the phylogenetic trees with the taxonomy system of these prokaryotes. It was a frequently asked question that what is the best length of the k-string, i.e., how to set the parameter *k*. As our studies, a reasonable length was in the range log_*m*_ *N* < *k* < log_*m*_*N* + 2 for the Hao method, where *m* is the number of letters in the type of genome and *N* is the average length of the genome sequences. And the reasonable *k* for new methods in the CVTree software should be a little bigger than that of the Hao method and have a larger value range. A detailed discussion of this problem can refer to our previous works [28, 29]. In this study, we set *k* = 6 for the Hao method, and *k* = 7 for the InterList method and the InterSet method. Figure 4 showed the relative entropy difference between the taxonomy and the phylogenetic trees at every taxon level. The relative entropy difference between the taxonomy and the phylogenetic tree from the LTPs132, which was obtained by the alignment method, was also plotted in the figure as the benchmark. It was found that the result of CVTree methods and LTPs132 have a similar performance at the high taxon levels of phylum, class, and order. At the low taxon level of family, genus, and species, however, the result of CVTree was more consistent with the taxonomy than that of LTPs132. That is, the taxa were more monophyly in the CVTree method. Moreover, our previous study showed that the CVTree methods may have much better performance with whole genomes for prokaryotes [30]. This indicated that the CVTree was an effective tool for conjecturing the taxonomy system.

**Figure 4.**
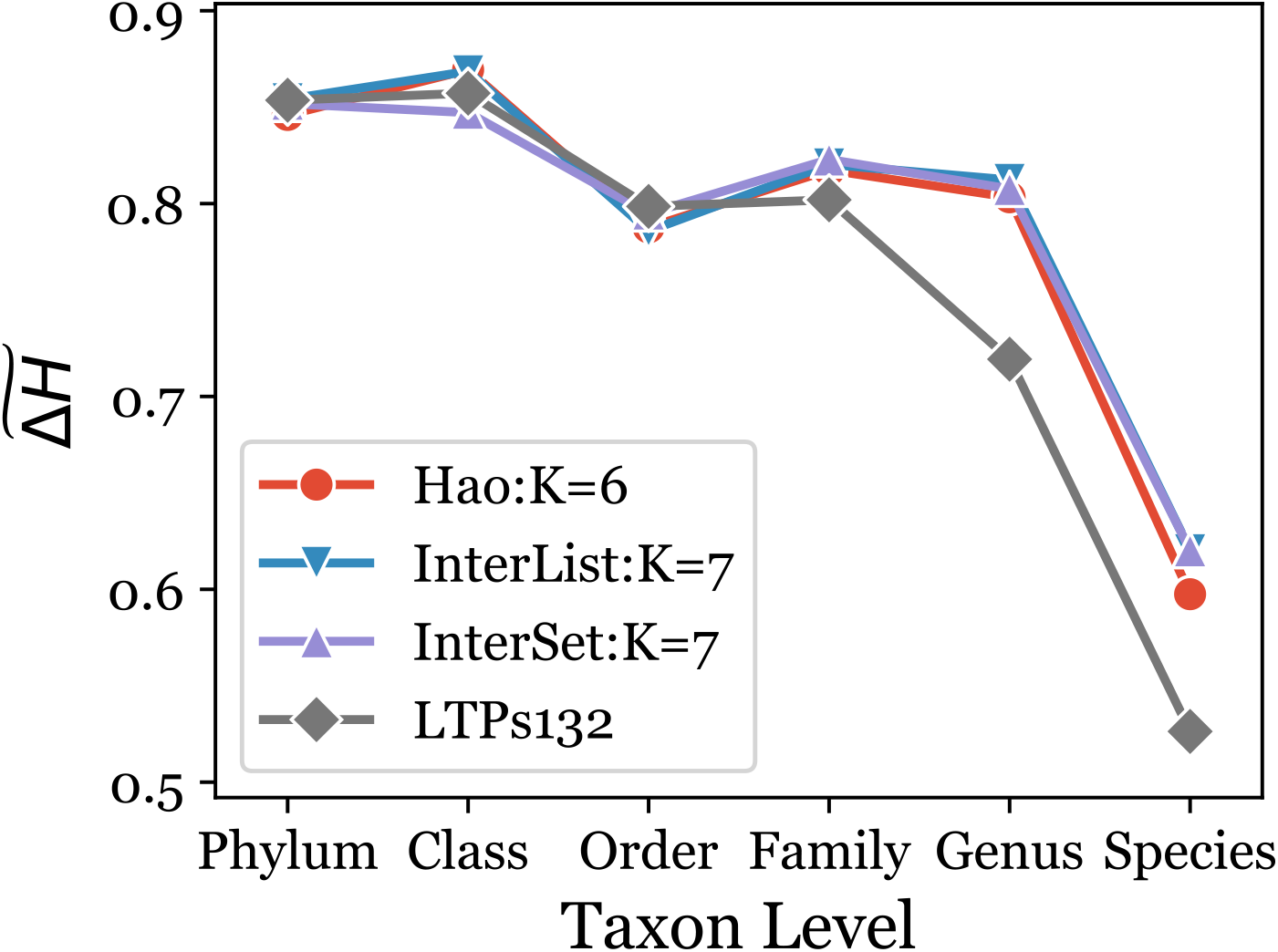
Relative Entropy Difference between Phylogeny and Taxonomy. Here the relative entropy difference 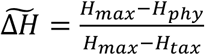, where *H* = − ∑_*i*_ *p*_*i*_ · log *p* is the Shannon entropy of a distribution. *H*_*phy*_ and *H*_*tax*,_ are the Shannon entropy of the distribution in phylogeny and taxonomy at a taxon level respectively. And *H*_*max*,_ indicated that every class including only one strain, i.e., *p*_*i*_ = 1/*N*. It is obvious that all taxa of a level make a partition of all strains, and *p*_*i*_ = *n*_*i*_/*N*. To obtain *H*_*phy*_ at a taxon level, we obtained the branches of all taxa of this level in the phylogenetic tree, it is also a partition of all strains. Thus, 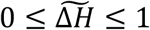, in which one indicated all taxa of the taxon level are monophyly, and 0 indicated every stain of all taxa are polyphyly in the phylogenetic tree.

## Conclusion

CVTree is a cluster of alignment-free methods to infer phylogenetic relationships based on genome sequences. It has been applied to viruses, prokaryotes, and fungi with remarkable success, as well as chloroplasts, mitochondria, and metagenomes. Here we released the standalone CVTree software. The main programs of the software were parallel by OpenMP techniques. It is efficient to obtain a taxonomy-compactible phylogenetic tree based on the 16S rRNA sequences. And since the complexity of the CVTree algorithm is lower than linear complexity with the length of genome sequences, CVTree is efficient to handle huge whole genomes, and obtained the phylogenetic relationship, especially for the prokaryotes. We believed that CVTree software may become an efficient and effective tool for establishing a phylogeny-based prokaryotic taxonomy.

## Author Contributions

GZ designed the study, wrote the source code, performed the analysis, and wrote the manuscript.

## Acknowledgments

We thank Dr. Qiang Li for his useful discussion. This work was supported by the National Basic Research Program of Ministry of Science and Technology of China (973 Project Grant No. 2013CB834100) and the National Natural Science Foundation of China (Grant No. 11474068). Author thanks the State Key Laboratory of Applied Surface Physics and the Department of Physics, Fudan University, China.

## References

[1] Needleman SB, Wunsch CD. A general method applicable to the search for similarities in the amino acid sequence of two proteins. J Mol Biol 1970;48:443–53.

[2] Smith TF, Waterman MS. Identification of common molecular subsequences. J Mol Biol 1981;147:195–7.

[3] Altschul SF, Gish W, Miller W, Myers EW, Lipman DJ. Basic Local Alignment Search Tool. Journal of Molecular Biology 1990;215:403–10.

[4] Thompson JD, Higgins DG, Gibson TJ. CLUSTAL W: improving the sensitivity of progressive multiple sequence alignment through sequence weighting, position-specific gap penalties and weight matrix choice. Nucleic Acids Res 1994;22:4673–80.

[5] Earl D, Nguyen N, Hickey G, Harris RS, Fitzgerald S, Beal K, et al. Alignathon: a competitive assessment of whole-genome alignment methods. Genome Res 2014;24:2077–89.

[6] Zielezinski A, Vinga S, Almeida J, Karlowski WM. Alignment-free sequence comparison: benefits, applications, and tools. Genome Biol 2017;18:186.

[7] Ren J, Bai X, Lu YY, Tang KJ, Wang Y, Reinert G, et al. Alignment-Free Sequence Analysis and Applications. Annual Review of Biomedical Data Science, Vol 1 2018;1:93–114.

[8] Zielezinski A, Girgis HZ, Bernard G, Leimeister CA, Tang KJ, Dencker T, et al. Benchmarking of alignment-free sequence comparison methods. Genome Biology 2019;20.

[9] Vinga S. Information theory applications for biological sequence analysis. Briefings in Bioinformatics 2014;15:376–89.

[10] Vinga S, Almeida J. Alignment-free sequence comparison - a review. Bioinformatics 2003;19:513–23.

[11] Qi J, Wang B, Hao B. Whole proteome prokaryote phylogeny without sequence alignment: a K-string composition approach. J Mol Evol 2004;58:1–11.

[12] Zuo G, Xu Z, Hao B. Phylogeny and Taxonomy of Archaea: A Comparison of the Whole-Genome-Based CVTree Approach with 16S rRNA Sequence Analysis. Life (Basel) 2015;5:949–68.

[13] Zuo G, Xu Z, Hao B. Shigella strains are not clones of Escherichia coli but sister species in the genus Escherichia. Genomics Proteomics Bioinformatics 2013;11:61–5.

[14] Zuo G, Hao B, Staley JT. Geographic divergence of “Sulfolobus islandicus” strains assessed by genomic analyses including electronic DNA hybridization confirms they are geovars. Antonie Van Leeuwenhoek 2014;105:431–5.

[15] Wang H, Xu Z, Gao L, Hao B. A fungal phylogeny based on 82 complete genomes using the composition vector method. BMC Evol Biol 2009;9:195.

[16] Kjaerbolling I, Vesth TC, Frisvad JC, Nybo JL, Theobald S, Kuo A, et al. Linking secondary metabolites to gene clusters through genome sequencing of six diverse Aspergillus species. Proc Natl Acad Sci U S A 2018;115:E753–E61.

[17] Gao L, Qi J. Whole genome molecular phylogeny of large dsDNA viruses using composition vector method. BMC Evol Biol 2007;7:41.

[18] Chu KH, Qi J, Yu ZG, Anh V. Origin and phylogeny of chloroplasts revealed by a simple correlation analysis of complete genomes. Mol Biol Evol 2004;21:200–6.

[19] Yuan J, Zhu Q, Liu B. Phylogenetic and biological significance of evolutionary elements from metazoan mitochondrial genomes. PLoS One 2014;9:e84330.

[20] Zhang Q, Wu Y, Wang J, Wu G, Long W, Xue Z, et al. Accelerated dysbiosis of gut microbiota during aggravation of DSS-induced colitis by a butyrate-producing bacterium. Sci Rep 2016;6:27572.

[21] Liu J, Wang H, Yang H, Zhang Y, Wang J, Zhao F, et al. Composition-based classification of short metagenomic sequences elucidates the landscapes of taxonomic and functional enrichment of microorganisms. Nucleic Acids Res 2013;41:e3.

[22] Qi J, Luo H, Hao B. CVTree: a phylogenetic tree reconstruction tool based on whole genomes. Nucleic Acids Res 2004;32:W45–7.

[23] Xu Z, Hao B. CVTree update: a newly designed phylogenetic study platform using composition vectors and whole genomes. Nucleic Acids Res 2009;37:W174–8.

[24] Zuo G, Hao B. CVTree3 Web Server for Whole-genome-based and Alignment-free Prokaryotic Phylogeny and Taxonomy. Genomics Proteomics Bioinformatics 2015;13:21–31.

[25] Li Q (2009), ‘A heuristic probabilistic model for the evolution of K-string of biological sequences and the problem of unique reconstruction of a sequence from its constituent K-string’, Department of Physics, Fudan University.

[26] Yarza P, Richter M, Peplies J, Euzeby J, Amann R, Schleifer KH, et al. The All-Species Living Tree project: a 16S rRNA-based phylogenetic tree of all sequenced type strains. Syst Appl Microbiol 2008;31:241–50.

[27] Yu J. A Scientist Guerilla Fighter in the Frontiers of Bioinformatics-In Memory of Bailin Hao. Genomics Proteomics – Bioinformatics 2018;16:307–9.

[28] Zuo G, Xu Z, Yu H, Hao B. Jackknife and bootstrap tests of the composition vector trees. Genomics Proteomics Bioinformatics 2010;8:262–7.

[29] Zuo G, Li Q, Hao B. On K-peptide length in composition vector phylogeny of prokaryotes. Comput Biol Chem 2014;53 Pt A:166–73.

[30] Zuo G, Qi J, Hao B. Polyphyly in 16S rRNA-based LVTree Versus Monophyly in Whole-genome-based CVTree. Genomics Proteomics Bioinformatics 2018;16:310–9.

